# Floristic, structural, successional and phylogenetic disparities in a mosaic of adjacent forest physiognomies in the Atlantic Forest

**DOI:** 10.64898/2025.12.19.695400

**Authors:** Yacov Kilsztajn, Heitor Lisboa da Fonseca, Joanna Marques Batista, Gláucia Cortez Ramos de Paula, Frederico Alexandre Roccia Dal Pozzo Arzolla

## Abstract

Tropical forests are structured as complex mosaics shaped by disturbance regimes and edaphic filters, yet distinguishing between successional dynamics and edaphic climax vegetation remains a challenge for forest ecology and conservation. This study examines the floristic composition, phytosociological attributes, successional stage, and phylogenetic diversity of three adjacent forest environments within Cantareira State Park, southeastern Brazil, one of the largest remnants of Atlantic Forest. Across 0.525 ha, we sampled 768 individuals, representing 92 species, 65 genera, and 42 families. Species richness was highest in environments B and C (56 species each) and lowest in A (42 species). Tree ferns (*Cyathea phalerata* and *Dicksonia sellowiana*) dominated the waterlogged environment A, while environment B was characterized by early secondary canopy trees and shade-tolerant understory species, and environment C by late secondary species such as *Ocotea catharinensis* and *Micropholis crassipedicellata*. Clustering analyses integrating floristic, structural, successional, and phylogenetic data revealed that environment C was consistently distinct, whereas A and B showed partial overlap due to shared edaphic conditions. Successional classification indicated apparent differences among environments, but the presence of mature forest indicators, including well-developed tree ferns and late secondary species, suggests that all three environments represent stable, mature forests shaped by contrasting soil and drainage conditions. These findings highlight the importance of recognizing edaphic climax communities within forest mosaics and caution against relying solely on successional stage classifications in conservation planning. Misinterpretation of structurally simple yet mature forests may bias management decisions and reduce protection under current legislation.

## 1. Introduction

Tropical forests are among the most biodiverse ecosystems on the planet (Barlow et al., 2018), but they are far from homogeneous (Whitmore, 1975, 1982; Hartshorn, 1980; Oldeman, 1983).

Instead, they are structured as complex mosaics of different vegetation types, each shaped by a variety of biotic and abiotic factors (Watt, 1947). These forest mosaics often emerge from two primary sources: natural disturbance regimes, such as gap dynamics (Carvalho, 1997; Whitmore, 1990; Pickett, 1983), and changes in local environmental conditions, especially edaphic variations (Clements, 1916 apud Oliveira e Silva Junior, 2011; Tansley, 1935 apud Oliveira e Silva Junior, 2011). This structural and compositional heterogeneity plays a fundamental role in forest dynamics, resilience, and long-term biodiversity maintenance (Clark et al., 1993; Balderrama and Chazdon, 2005).

In mature forests, environmental filters such as soil type and drainage conditions can give rise to edaphic climax communities, vegetation types that persist under specific soil or hydrological constraints, independently of disturbance history (Hulshof and Spasojevic, 2020). These communities may host species typically associated with early successional stages, not because of recent disturbance, but due to physiological adaptations to limiting conditions such as poor nutrient availability, shallow soils, or prolonged waterlogging (Pan et al., 2021; Xavier et al, 2019).

In flood-prone or poorly drained areas, water saturation can significantly limit oxygen availability in the root zone, restricting the growth of many plant species and favoring those with specific ecological strategies for coping with hypoxia (Pan et al., 2021). These conditions result in vegetation with distinct structural and floristic features, and different species assemblages compared to well-drained upland forests (Clark et al., 1999). Importantly, such environments can be misinterpreted as degraded or regenerating forests when, in fact, they represent mature, stable systems adapted to edaphic constraints.

This distinction has profound implications for conservation and forest management, particularly in tropical biodiversity hotspots like the Atlantic Forest. As a critically endangered biome, the Atlantic Forest has suffered extensive deforestation, fragmentation, and degradation, with less than 15% of its original cover remaining (Ribeiro et al., 2009). Efforts to conserve and restore this biome depend on accurate assessments of forest condition and successional stage (Resende et al., 2024). However, the presence of structurally simple or floristically distinct patches within mature forest mosaics can lead to misclassification of their conservation value, potentially opening the door to legal exploitation or insufficient protection.

In Brazil, environmental legislation often links forest use restrictions to the successional stage of vegetation (Resende et al., 2024). Areas classified as early-stage regeneration may be subjected to less stringent conservation measures or even legal logging, whereas mature forests receive higher levels of protection. In this context, understanding the true nature of forest mosaics, including the role of edaphic and hydrological constraints, is essential for avoiding misinterpretation and ensuring appropriate conservation strategies.

In this study, we investigate the ecological complexity of a forest mosaic within Cantareira State Park (Parque Estadual da Cantareira – PEC), one of the largest urban tropical forests in the world and a key remnant of the Atlantic Forest in southeastern Brazil (Mattos et al., 2010). We focus on three adjacent forest physiognomies that differ in soil and drainage conditions but occur within the same mature forest matrix. Specifically, we ask: (1) Are the forests developed over these distinct soil environments different from each other in terms of floristic composition, phytosociological structure, and phylogenetic structure? and (2) Do these differences translate into variation in the successional classification of the species that compose them? We hypothesize that the three forest types are indeed distinct in their floristic, structural, and phylogenetic composition due to the strong influence of edaphic filters. Furthermore, although all sites represent mature forests, we expect the successional classification of their species to vary according to soil drainage conditions—forests established on more waterlogged soils are expected to exhibit a simpler apparent successional structure, with a higher proportion of species typically associated with early or intermediate successional stages.

## 2. Materials and Methods

### 2.2 Study site

This study was conducted in Cantareira State Park (Parque Estadual da Cantareira – PEC), which covers 7,916.52 ha in the Serra da Cantareira, Upper Tietê River Basin, Atlantic Plateau, Brazil. The park ranges from 875 to 1,215 m in elevation and features rugged terrain with steep slopes and incised valleys (São Paulo, 2010). The climate is humid temperate without a dry season (Cfb, Köppen), and vegetation is mainly Montane Dense Ombrophilous Forest, composed largely of secondary Atlantic Forest with remnants of primary forest (IBGE, 2012; Arzolla, 2002; São Paulo, 2010).

The study was carried out in the Pinheirinho Region (Figure S1), within the Ribeirão São Pedro micro-basin, one of the best-preserved areas of the park (Baitello et al., 1993). The landscape includes hills, mountains, and seasonally waterlogged plains, forming a mosaic of forest types that increases ecological complexity (São Paulo, 2010). We selected three adjacent forest environments with contrasting conditions (environments A, B, and C). The three environments are 100–150 m apart, within a total study area of ∼1.5 ha. Environments A and B occur on low-slope terrain with alluvial forest: A is permanently waterlogged on Melanic Gleysol, while B lies on Humic Latosol, near streams but not waterlogged; in these environments, the riverbed causes canopy fails and consequently greater light entry. Environment C is a north-facing hillside with shallow, well-drained Haplic Cambisol and slope forest. The soils of the three environments were characterized by auger drilling to 1 m depth, with composite samples collected from 0–20 cm and 40–60 cm.

### 2.3 Field sampling

In each environment, we established seven plots measuring 10 × 25 m, totaling 0.525 ha of sampled area. We included all living trees and arboreal ferns with a stem perimeter at breast height (PBH, measured at 1.30 m above ground) ≥ 15 cm. We tagged each individual with a numbered metal label for identification. Whenever possible, we identified species directly in the field. For individuals requiring confirmation, we collected botanical material, which we later identified through comparison with herbarium specimens, consultation with taxonomic specialists, and specialized literature.

Taxonomic nomenclature follows the Flora e Funga do Brasil (2020) and Angiosperm Phylogeny Group IV (APG, 2016).

### 2.4 Forest structure and diversity

In each environment, we characterized the floristic and successional composition, forest structure, and phylogenetic diversity. We determined the successional composition by classifying species into four groups, following previous studies (Bernacci et al., 2006; Catharino et al., 2006; Arzolla et al., 2020; Silva et al., 2022). Pioneer species are light-demanding and occur mainly in edges and large canopy gaps; early secondary species grow in subcanopies or small gaps with moderate light; late secondary species establish in shaded understories and may reach the upper stratum or the canopy of the forest; and shade-tolerant or typical of understorey species complete their life cycle in deep shade.

We evaluated forest structure using standard phytosociological parameters, including relative density (DeR), dominance (DoR) and frequency (FR), following Mueller-Dombois & Ellenberg (1974). All analyses were performed in Fitopac 2.1 (Shepherd, 2010). We quantified phylogenetic diversity using Faith’s Phylogenetic Diversity (PD), and accounted for species richness by comparing observed values against null models generated from 999 randomizations. To reconstruct phylogenetic relationships among species, we used the U.PhyloMaker R package (Jin and Qian, 2023), and we calculated diversity metrics with the picante R package (Kembel et al., 2010).

### 2.5 Comparisons among environments

To compare environments, we conducted multivariate cluster analyses of all sampled plots using the Unweighted Pair Group Method with Arithmetic Mean (UPGMA) based on the Bray–Curtis coefficient, implemented in the vegan R package (Oksanen et al., 2013). These analyses allowed us to group plots according to overall similarities, highlighting patterns of ecological resemblance among sites. Floristic composition disparities were assessed using species abundance matrices, providing insight into differences in species presence and relative dominance across plots. Successional structure was evaluated by examining species abundances per successional class, which helps reveal how ecological succession stages vary among environments. Structural differences were quantified using species basal area matrices, capturing variation in vegetation architecture and biomass distribution.

Finally, phylogenetic disparities were estimated using UniFrac distances, implemented in the rbiom R package (Smith, 2021), providing a measure of evolutionary differences among communities that integrates both species presence and their phylogenetic relationships.

## 3. Results

### 3.1 Forest structure and diversity

In total, we sampled 768 individuals, representing 92 species, 65 genera, and 42 families (Table 1; see Table S1 for details). Species richness was highest in environments B and C (56 species each) and lowest in environment A (42 species, ∼25% fewer). Across all plots, the richest families were Myrtaceae (16 species), Lauraceae (eight), and Rubiaceae (seven), while the most abundant were Rubiaceae (106 individuals), Cyatheaceae (103), Myrtaceae (65), and Fabaceae (36). Regarding community parameters (Table 1), environment A consistently showed the lowest richness, diversity (SW and PD), dominance, and density, likely due to the constraints of its waterlogged soils. Environments B and C were similar in most metrics, although dominance was 20% higher in B (Table 1), reflecting a greater abundance of large individuals with higher basal area.

**Table 1.** Diversity and structure metrics for each sampled forest environment.

| Summary metrics | Total values | A | B | C |
| --- | --- | --- | --- | --- |
| Species richness | 93 | 42 | 56 | 56 |
| Density (ind./ha) | 1462.86 | 1011.43 | 1685.71 | 1691.43 |
| Dominance (m<sup>2</sup>/ha) | 46.2 | 33.76 | 57.92 | 46.37 |
| Shannon-Wiener index (SW) | 3.961 | 3.123 | 3.454 | 3.465 |
| Phylogenetic diversity (PD) | - | 3648.809 | 4506.664 | 4468.181 |
| Standardized PD | - | 0.316 | 0.972 | 1.341 |

In environment A, the most abundant families were Cyatheaceae and Dicksoniaceae, while Myrtaceae and Euphorbiaceae stood out in species richness (Table S1). Density and dominance were concentrated in the tree ferns Cyathea phalerata and Dicksonia sellowiana (Figure 1, Table S2), reflecting their adaptation to waterlogged alluvial soils. In environment B, the most abundant families were Cyatheaceae and Rubiaceae, with Myrtaceae and Lauraceae richest in species (Table S1). The most abundant species were *Psychotria suterella* and *Alsophila setosa*, while relative dominance was led by *Cinnamomum pseudoglaziovii, Cedrela fissilis*, and *Alchornea triplinervia* (Figure 1, Table S3). Except for *C. pseudoglaziovii* (late secondary), this environment was dominated by large early secondary canopy trees and shade-tolerant understory species.

**Figure 1.**
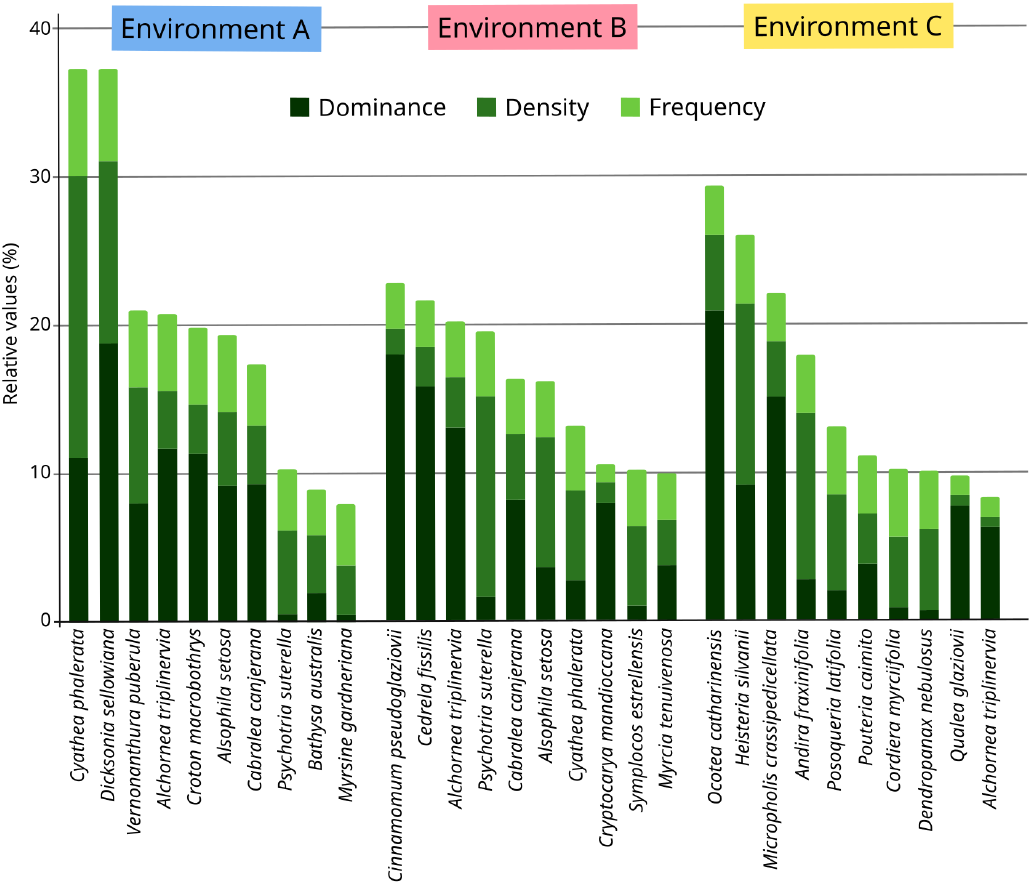
Relative dominance, density, and frequency of the ten species with the highest importance values in each environment.

In environment C, the most abundant families were Rubiaceae, Erythropalaceae, and Fabaceae, while Myrtaceae and Rubiaceae were richest in species (Table S1). The most abundant species were *Heisteria silvianii, Andira fraxinifolia, Posoqueria latifolia*, and *Dendropanax nebulosus* (Figure 1, Table S4). Dominance was concentrated in *Ocotea catharinensis, Micropholis crassipedicellata, H. silvianii*, and *Qualea glaziovii*, all late secondary species, indicating a more advanced successional stage than in the other environments.

### 3.2 Comparisons among environments

Although the three environments are located very close to each other, clear differences emerge in floristic, structural, successional, and phylogenetic composition when we examine the clusters formed by the sampled plots (Figure 2). Environment C is consistently distinct from environments A and B across all dimensions, whereas the distinction between A and B is less pronounced, which is expected given that both are alluvial environments. At the structural and successional levels, the edaphoclimatic similarities shared by A and B, both occurring in low-slope terrain with alluvial forest, likely reduce the degree of separation between them. Interestingly, when we include the phylogenetic component, the plots in environment B are more similar to each other than to those in environment A, while the reverse is not true. This suggests that part of the phylogenetic diversity present in environment A is also represented in environment B.

**Figure 2.**
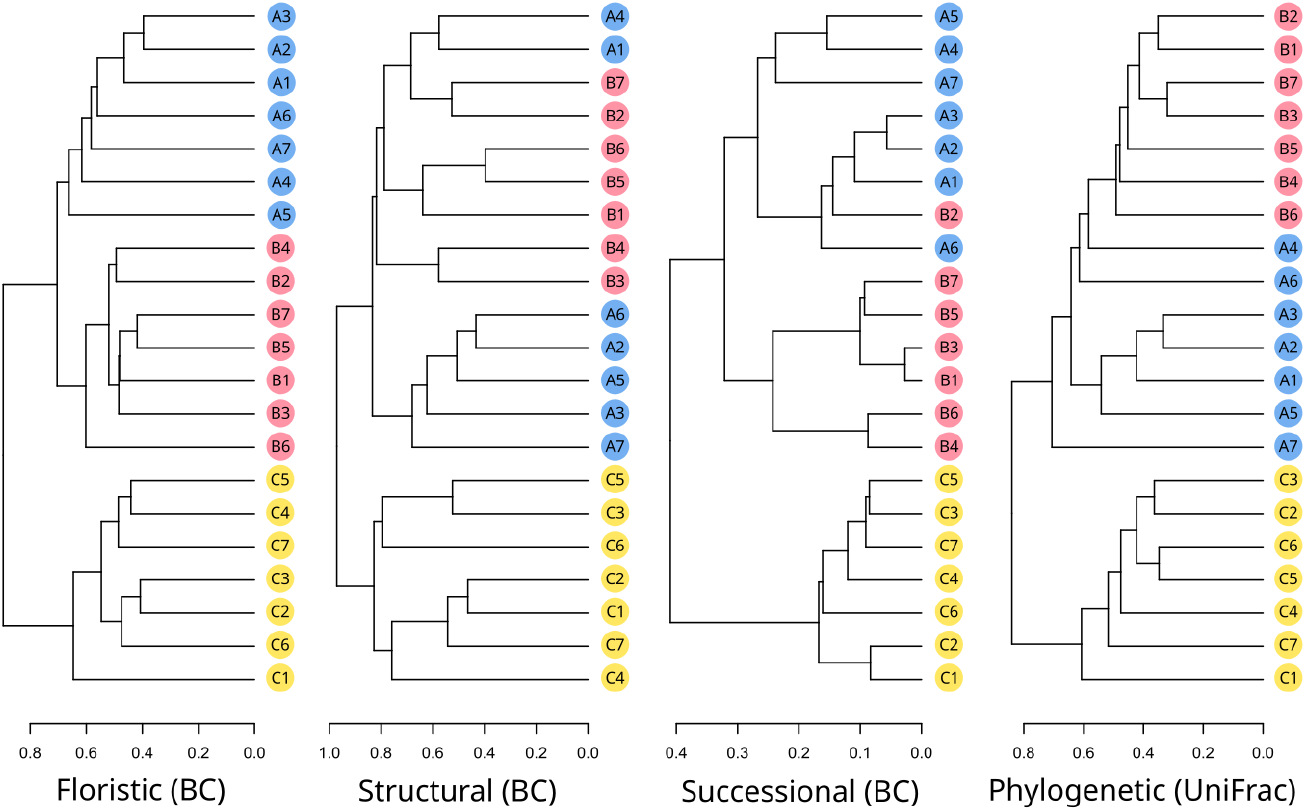
Floristic, structural, successional, and phylogenetic disparities among sampled plots. Dendrogram tip codes indicate environment type and plot number. Clustering was based on the Bray–Curtis coefficient (BC) and UniFrac distances.

Regarding the successional classification of the three environments, both the percentage of species and of individuals (Figure 3 and Table S1) in each successional class reveal that they differ in composition and structure. In environments A and B, shade-tolerant, pioneer, and early secondary species predominate, with pioneers more common in A and early secondaries more frequent in B. Environment C, in contrast, is characterized by the dominance of late secondary species (Figure 3).

**Figure 3.**
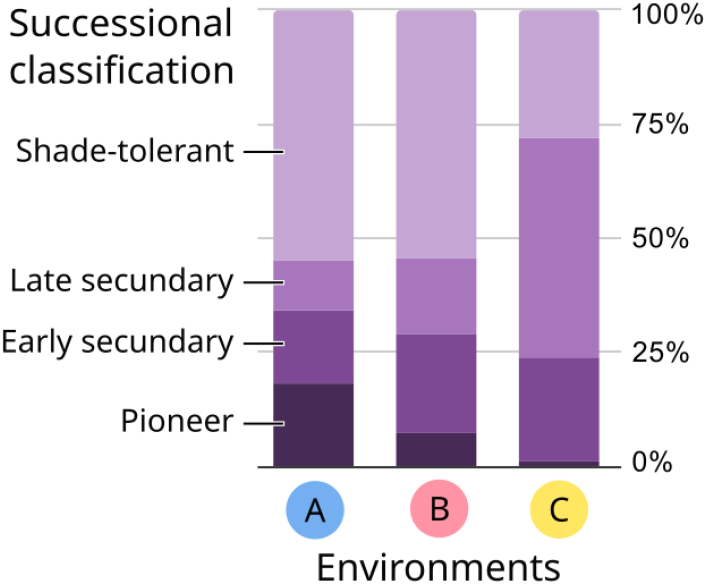
Successional classification of individuals per environment, expressed as the percentage of individuals in each successional class.

Based on the historical context of the area and the presence of key indicators — such as well-developed tree ferns and late secondary species in environments A and B (Table S1) — it is evident that all three environments represent mature forests, even though the overall pattern suggests distinct successional stages. The differences observed in successional species classification are most likely related to contrasting soil conditions, resulting in distinct edaphic climaxes in each environment. These patterns underscore potential pitfalls: assessments of forest successional stages may lead to biased management and conservation decisions if the existence of multiple structural configurations within mature forests is not considered.

## 4. Discussion

The composition and structure observed across the three forest environments reflect well-documented environmental gradients in the Atlantic Forest domain. Environment A, dominated by Cyatheaceae and Dicksoniaceae and featuring lower richness and density, resembles permanently water-logged alluvial forests, where tree ferns and hydromorphic-soil adapted taxa tend to dominate (Scarano 2002; 2009). Environments B and C, while similar in richness, differ in structure: B shows greater dominance linked to large individuals and intermediate-successional species, whereas C is characterised by late-successional taxa, a pattern consistent with successional and hydrological variation reported in other alluvial/slope forests of the Atlantic Forest (Silva et al. 2020). These parallels underscore that local factors (edaphic conditions, topography situations and hydrological regime) are crucial in determining beta-diversity among adjacent plots. Finally, our findings concerning the relative importance of Myrtaceae, Lauraceae and Rubiaceae, as well as the structural role of both arboreal stems and tree ferns, align with floristic inventories of alluvial Atlantic Forests and highlight the need for management approaches that account for environmental heterogeneity in local conservation and restoration planning (Silva et al. 2020, Scarano 2002).

Adopting a multidimensional perspective to assess community differentiation provides a more comprehensive understanding of forest heterogeneity than analyses based solely on species composition or structure. Traditional floristic or structural comparisons often capture only part of the ecological variation among environments, potentially overlooking evolutionary or functional processes that shape assemblage composition (Devictor et al. 2010, Graham and Fine 2008). By jointly evaluating floristic, phytosociological, successional, and phylogenetic dimensions, the interpretation of divergence among sampled areas becomes more robust (Webb et al. 2002, Cavender-Bares et al. 2009). In our case, we observed that the degree of separation between environments A and B varied depending on the dimension considered, suggesting that these two environments share certain ecological affinities, likely linked to their similar edaphoclimatic conditions. In the Atlantic Forest, few studies have simultaneously examined these complementary dimensions; however, recent research has shown that incorporating phylogenetic information can reveal hidden layers of beta diversity and successional dynamics even across short spatial gradients (Gastauer et al. 2017, Dvorak et al. 2025).

This multidimensional framework therefore enhances our ability to interpret community patterns in environmentally complex landscapes such as the Serra da Cantareira, where fine-scale heterogeneity and disturbance history strongly influence both species turnover and lineage composition.

Our observations resonate with the concerns raised by Resende et al. (2024), who emphasize that classifying Atlantic Forest remnants into discrete successional stages can oversimplify complex ecological trajectories. This categorical approach assumes a single, linear pathway of forest development, disregarding the influence of local filters such as soil, hydrology, topography and disturbance history that generate multiple, stable structural configurations. In this context, the distinct combinations of species and structural attributes observed among environments A, B, and C are better understood as expressions of alternative mature states rather than sequential successional phases. As highlighted by Resende et al. (2024), such misinterpretations may have significant conservation consequences, since management and restoration programs that rely on rigid successional labels risk undervaluing forests that are ecologically mature but structurally divergent. Adopting a more flexible, process-oriented framework that recognizes these alternative pathways is therefore essential for accurately assessing forest integrity and ensuring that conservation priorities reflect true ecological resilience.

## 5. Conclusion

Our results demonstrate that the three forest physiognomies within Cantareira State Park differ significantly in floristic composition, phytosociological structure, phylogenetic organization, and successional composition, supporting our initial hypothesis that edaphic conditions play a major role in shaping local forest diversity. Although all sites represent mature forests, variations in soil type and drainage lead to distinct community assemblages and successional profiles. Forests established on poorly drained, waterlogged soils show simpler structural and successional patterns, dominated by species tolerant to hypoxia and nutrient limitations, while well-drained areas sustain higher diversity and more complex canopy organization. These findings highlight that physiognomic simplicity in such environments does not necessarily indicate regeneration or degradation but can represent stable, edaphically constrained climax communities. Recognizing these distinctions is crucial for accurate forest classification, avoiding misinterpretations in conservation assessments, and ensuring that mature forests adapted to extreme edaphic conditions receive adequate legal protection and management attention.

## 6. Acknowledgements

We thank the Instituto de Pesquisas Ambientais for supporting this research and the National Council for Scientific and Technological Development (CNPq) for grants awarded to Y. Kilsztajn, J.M. Batista, and H.F. Lisboa, under the supervision of F.A.R.D.P. Arzolla. We are grateful to J.B. Baitello and O.T. Aguiar for identifying Lauraceae and Myrtaceae species, respectively; to M. Rossi for soil identification; and to M.M. Kanashiro for valuable insights regarding Figure S1.

## 7. Author contributions (CRediT)

Conceptualization and Methodology: FARDP; Data curation: all authors; Formal analysis and Visualization: YK; Funding acquisition and Supervision: FARDP; Writing – original draft: HLF and YK; Writing – review and editing: all authors.

